# Morphometric analyses of shape: The analysis software toolbox for quantification of craniofacial shape

**DOI:** 10.1101/2025.11.14.688515

**Authors:** Makenzie C. Stearsman, Jennan A. Lahamer, Raèden Gray, C. Ben Lovely

## Abstract

Fetal alcohol spectrum disorders (FASD) are characterized by a varying set of physical, cognitive, and behavioral disabilities caused by prenatal ethanol exposure, including those to the facial skeleton. Ethanol-sensitive genetic loci contribute to this high degree of variation in FASD, which complicates analyses of facial shape. We have previously shown that we can analyze these changes in facial shape from gene-ethanol interactions in zebrafish. Zebrafish are an ideal model to analyze this variation for several reasons; 1. 70% of genes are orthologs between humans and zebrafish; 2. external fertilization allows researchers to control timing and dosage of ethanol treatments; 3. the structure of their facial skeletal is conserved with vertebrates; 4. translucent larvae enable direct viewing of changes to the craniofacial skeletal structure during development. However, analyzing the shape of the craniofacial skeleton can be difficult to fully assess through simple linear measures, as these do not capture overall changes in shape. In addition, changes in head size can complicate data interpretation. To address this, we undertook a morphometric approach, analyzing overall facial shape through principal component analyses via freeware software, TPSDigs2, MorphoJ, and PAST. The combination of this software allows for pairwise comparison of overall facial morphology. Here, we outline our approach and analysis of facial shape in ethanol-treated zebrafish mutants using these programs to conduct a series of complementary multivariate statistical analyses.

**SUMMARY:** This work describes a protocol for quantifying craniofacial cartilage shape using free software (TPSDigs2, MorphoJ, and PAST) to measure changes in facial structure in zebrafish larvae.

## INTRODUCTION

Fetal Alcohol Spectrum Disorders (FASD) are characterized by a broad range of developmental defects, including behavioral, neurological, and physical [1, 2]. Included in these physical defects are craniofacial defects, such as jaw hypoplasia [3-6]. While timing and dosage of ethanol exposure contribute to the complex etiology of FASD, genetic contribution plays a significant role in FASD etiology [6-12]. This combination of factors makes studying facial defects in human cohorts challenging. To study the impact of prenatal ethanol exposure on development, we use the zebrafish model. Zebrafish serve as a strong model for FASD as they share 70% gene orthologs with humans, 82% of known disease-causing genes [13-16]. In addition to genetic conservation, zebrafish: 1. have high fecundity, allowing for several zebrafish larvae to be produced at a time, 2. undergo external fertilization, which allows direct study of ethanol-sensitive developmental processes, 3. have highly conserved and stereotyped vertebrate craniofacial development, and 4. Are translucent embryos/larvae allowing for visualization of skeletal structures [17]. We have previously shown that mutation in many different zebrafish genes sensitizes embryos to ethanol-induced facial defects, and these defects can be subtle and difficult to identify by eye [10, 18]. In addition, we observed that ethanol-treated wild-type larvae also have slight changes in the craniofacial region compared to untreated wild-type larvae, though again these changes are difficult to identify by eye [10]. Although these changes go undetected upon visual observation, we were able to show these ethanol-induced changes in facial shape using morphometric analyses available on 2D images of the viscerocranium [10].

The face is a complex 3D structure that is difficult to measure using conventional linear measurements. These single linear measures do not account for the relative position of each structure measured to, and their impact on, the other measured structures of the face. Morphometric analyses address these shortcomings by using shape configurations via landmarks that account for the relative position of all structures measured, even controlling for overall differences in size [18-20]. This approach results in data that is more accurate, has greater resolution, and is much easier to visualize, which can yield results that may not be observed using conventional linear measurements [18-20]. In addition, morphometric approaches can make direct comparisons between and within groups using multivariate statistical methods. Here we describe our use of the freeware morphometric software, TPSUtil, TPSDigs2, MorphoJ, and Paleontological Statistics (PAST), in combination with 2D images of the viscerocranium in untreated and ethanol-treated wild-type and *bmp7a* heterozygous larvae. While these programs have been utilized mainly in paleontology and ecology-focused fields, when used in combination, they provide a thorough analysis of facial shape. Ultimately, in this protocol, we show how to use these freely available software applications to quantify craniofacial cartilage shape changes in the zebrafish model.

## PROTOCOL

### ANIMAL USAGE

All zebrafish larvae used in this procedure were raised and bred following established IACUC protocols [21]. These protocols were approved by the University of Louisville.

### ZEBRAFISH STRAINS USED

The zebrafish strain *bmp7a*^*ty68a*^ [22] and wildtype siblings (all in the AB background) were used in this study. All the water used in this procedure was sterile reverse osmosis water. Adult fish were maintained at 28.5°C with a 14 / 10-hour light / dark cycle. Sex as a biological variable does not apply to our studied development stages, as sex is first detectable in zebrafish around 20-25 days post-fertilization [23], after all of our analyses.

### ETHANOL TREATMENT

Eggs from random heterozygous crosses were collected, and embryos were morphologically staged [21], sorted into sample groups of 100, and reared at 28.5°C to desired developmental time points. All groups were incubated in embryo media (EM) [10, 18]. At 6 hours post fertilization (hpf), EM was changed to either fresh EM or EM containing 1% ethanol (v/v). At 24 hpf, EM containing ethanol was washed out with three fresh changes of EM [10, 18].

### FACIAL STAINING AND IMAGING

Zebrafish larvae were fixed at 5 days post fertilization (dpf) and stained with alcian blue, labeling cartilage [24]. Whole mount, ventral view, brightfield images of the viscerocranium were taken on an Olympus BX53 compound microscope as previously described [25].

### GENOTYPING

The tails from 5 dpf old larvae were removed from *bmp7a* larvae following Alcian Blue cartilage staining as previously described [18]. The heads and tails of the larvae were separated. The heads remained in glycerol for storage for later imaging. Tails were lysed with 5 µL Proteinase K (10 mg/mL – Millipore Sigma) to extract gDNA. PCR was then performed on the gDNA for each sample using primers targeting genes *bmp7a* to amplify regions containing the SNP in *bmp7a*. Restriction endonuclease-based DNA digestion using BslI (NEB) differentiates mutant from wild-type alleles based on the SNP. The samples were then divided by genotype into pooled groups and were imaged for analysis.

### SOFTWARE USED FOR SHAPE ANALYSES

Morphometric analysis of alcian-stained larvae was performed with TpsDig2 (https://sbmorphomectrics.org), MorphoJ [20] and Paleontological Statistics Software Package for Education and Data Analysis (PAST) analysis software [26]( [https://past.en.lo4d.com/windows]). Principal component analysis (PCA), Canonical Variate Analysis (CVA), Procrustes ANOVA, and wireframe graphs of facial variation were generated using MorphoJ. All the statistical analyses were performed using MorphoJ and PAST software [20, 26].

#### 1. Image the facial skeleton consistently for all samples to be analyzed

1.1 Images of the facial skeleton should be taken at identical microscope settings and saved as .tif files

1.2 Files should be saved in a single folder labeled to fully identify the experiment

#### 2. Preparing images with TPS software

2.1. Open Notepad and select “Save As”. From here, click “all files” at the bottom. Save the file as “experiment-name.tps” and save it in your folder containing all images to be analyzed.

#### 3. Inputting images into TPS software

3.1. Open TpsUtil software. Next, pull down “Operation” and select the “Build tps file from images” tab.

3.2. From this tab, click “input” and open the file containing your images. Click the first image in the file and select “open”.

3.3. Select the “output” tab, followed by “tps notepad file” and save. When prompted to replace the file, select “yes”.

3.4. Next, click the “set up” tab and “create”. Go to the file to see if the images have been added.

#### 4. Adding landmarks to images

4.1 Open TPSDigs2 and open the following tabs in order: “Input source”, “file”, then the saved “TPS Notepad file.”

4.2. Next, click “options” to visualize the “Image tools” option. Here, make sure to set the reference length to 100 microns. Press “set scale”.

NOTE: Do not scale relative to the scale on the image.

4.3. When setting the scale, press “ok” to set parameters, then exit image tools.

4.4. At this point, add landmarks to each image to be analyzed for proper morphology analysis by clicking the aim symbol.

4.5. Landmarks were placed on the following joints (See Fig. 1E):

**Figure 1.**
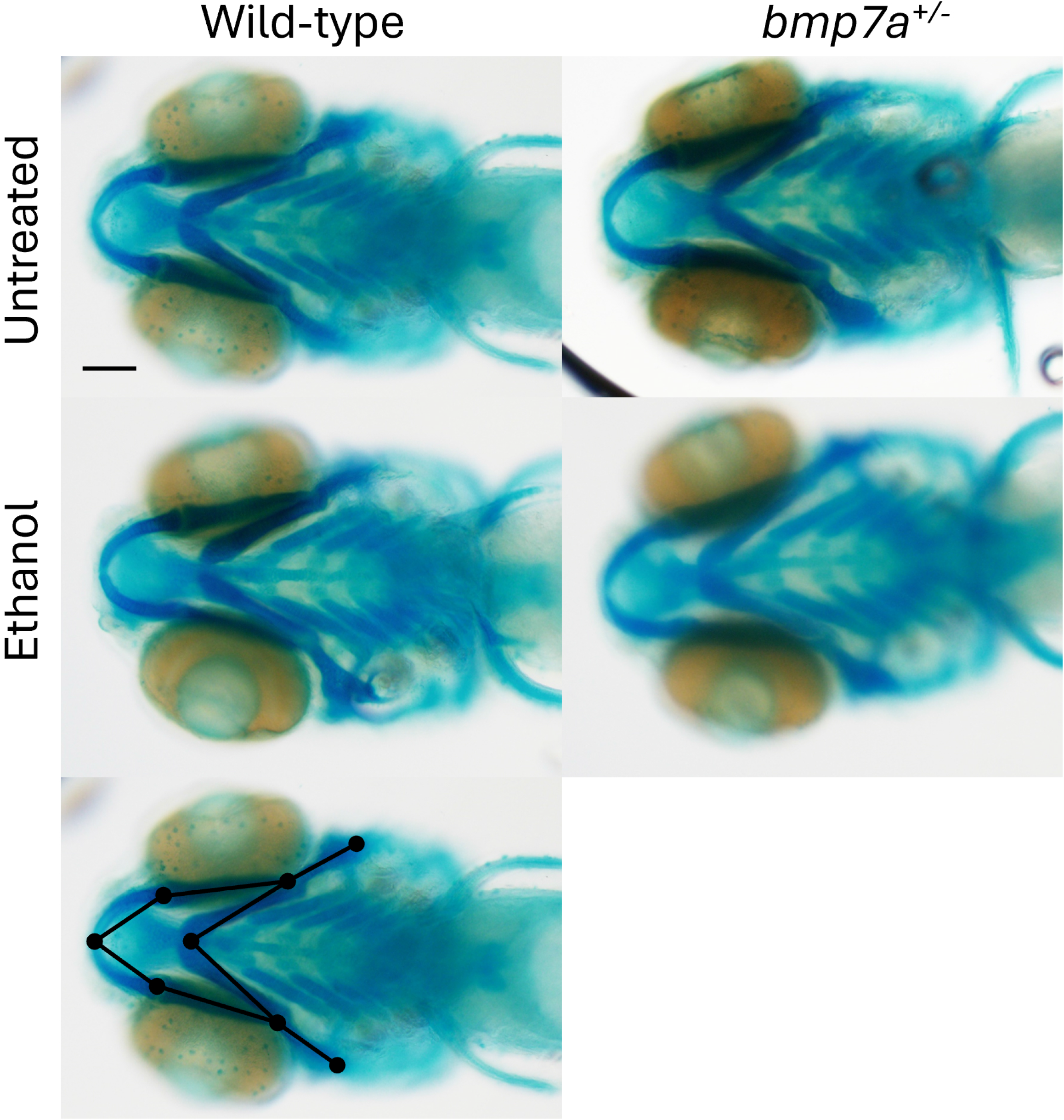
Alcian-stained zebrafish larvae. (A-E) Whole-mount images of viscerocranium at 5 dpf larva. Cartilage is blue (Ventral views, anterior to the left, scale bar: 100 μm). (A) Wild-type larvae untreated (n = 10). (B) Wild-type larvae treated with ethanol 6-24 hpf (n = 4). (C) bmp7a+/- untreated (n = 8). (D) bmp7a+/- treated with ethanol 6-24 hpf (n = 8). (E) Example landmark/wireframe on wild-type larva from panel (A).

4.5.1. Midline joint between the Meckel’s cartilages

4.5.2. The bilateral joints between Meckel’s and the palatoquadrate cartilages

4.5.3. Midline joint between the ceratohyal cartilages

4.5.4. The bilateral joints between the palatoquadrate and ceratohyal cartilages

4.5.5. The distal end of the hyomandibular cartilages.

4.6. After placing landmarks on an image, click “files” for the dropdown menu. Follow this pathway to “save data”, and then “overwrite”.

4.7. Exit TPSDigs2.

#### 5. Using MorphoJ to analyze shape by Procrustes ANOVA

5.1. Open MorphoJ software.

5.2. Under preliminaries, create and name the data set. Select “create new data set” and name the data set. Then, click “TPS” and select the notepad with the new data points added from step 4.4. Create the dataset.

5.3. The images are ready to be statistically viewed using Procrustes fit. Select the tab reading “project tree” and click on your data set. Select the following buttons in this pathway: “preliminaries”, “new Procrustes fit”, “Align by principal axes”, “Perform Procrustes fit.”

5.4. Select “generate covariance matrix” under preliminaries; the software will say Procrustes coordinates. Execute the function when prompted.

5.5. Select “create or edit wireframe” under preliminaries and link the points on the images. Then select “link points” and “accept” or “create” the image. Edit classifiers.

5.6. Open Microsoft Excel while keeping MorphoJ open. Add all necessary information, for example: Genotype, Treatment, Genotype and Treatment, Experiment, etc. Save the Excel sheet as a CSV file.

5.7. In MorphoJ, click “file” and “import classifier variables” to select the CSV file. After opening the file, go back to the project tree and click the dataset.

5.8. Once the dataset is opened, click “preliminaries” and “edit classifiers”. This will ensure all images are added.

5.9. Next, click “project tree”, “Covmatrix”, then “Variation” at the top.

5.10. Select “Principal component analysis” to see PC scores. Click “PC scores to see the generated graph.

5.11. To add colors to the graph, right-click and press “confidence ellipsis” to add the desired classifier. Press “color data points” to add your colors, and “ok” to accept these changes.

NOTE: Once you close out of the “PC score” tab and go back in following the instructions in step 5.10, the colors should be updated.

5.12. To change the wireframe colors, go to “Preliminaries”. Click “set options” for the shape graph, located at the bottom of the screen. Select “wireframe graphs” to change the colors of the target shape, starting shape, and numbers.

5.13. Select “Variation” then “Procrustes ANOVA”.

5.14. Export the Procrustes ANOVA results to the Microsoft Excel file from step 5.7.

#### 6. Canonical Variate Analysis of the dataset

6.1. In MorphoJ, select the original dataset or project. To begin Canonical Variate Analysis (CVA), select “Comparison” followed by “Canonical Variate Analysis”. Select the classifier variable(s) – select Genotype & Treatment and execute the function.

6.2. To export CVA, click the “results” tab and right-click on the now-open results page. Select “Export to File” and save the information.

6.3. Click Project Tree and select “CVA”, then “Scores”. At the top left of the page, select “File”. Once the File tab has opened, choose “export dataset” and select the data type and Genotype & Treatment. Save CVA scores as a .txt file.

6.4. To prepare the file for PAST software, open the saved CVA scores in Notepad or an equivalent software. Change “ID” in the top left corner to “Label” so as not to confuse PAST software. Save the edited CVA Scores or equivalent.

#### 7. MANOVA in PAST software

7.1. To import the CVA scores in PAST, select “File” the “Open” to access the saved CVA scores from Notepad or equivalent. When the “Import text file” window opens, select “Names, Data” for row and column and “tab” for a separator. Select “Import”.

7.2. Under the “show” tab, select “Column Attributes”. Next to the “Type” button, open the dropdown menu and select “Group for the first column that should have the classifier variable(s).

7.3. Click the gray empty cell in the top left corner, above “type”, to select the entire dataset. Select “Multivariate”, “Tests”, and “MANOVA” to process the MANOVA. Export the MANOVA results to the Microsoft Excel file from step 5.7.

NOTE: You will be given both “Summary” and “Pairwise” results. Export or save both.

## REPRESENTATIVE RESULTS

To identify ethanol-induced facial shape changes in zebrafish, we combined freely available software applications to generate and quantify morphological data of the facial skeleton. Embryos from *bmp7a* heterozygous adult carrier fish were generated and treated with ethanol from 6-24 hpf. Larvae from these crosses were fixed at 5 dpf, and facial cartilages were stained with Alcian Blue. Images of the viscerocranium were taken for each larva in each genotype and treatment group (Fig. 1A-D). Landmarks were placed on each image using TPSDigs2 and saved (Fig. 1E). This dataset of “landmark-labeled” images was then imported into MorphoJ. Principal component analysis (PCA) was conducted to visualize the distribution of individuals based on shape variation. Each Principal component (PC1, PC2, etc.) shows specific variation in viscerocranial shape (Fig. 2, magenta wireframe) relative to the average shape of all viscerocrania (Fig. 2, black wireframe). PC1 and PC2 accounted for the majority of total shape variance, with PC1 representing approximately 34% of all variation and PC2 representing 20% of all variation, reflecting subtle but continuous variation across all individuals (Fig. 3).

**Figure 2.**
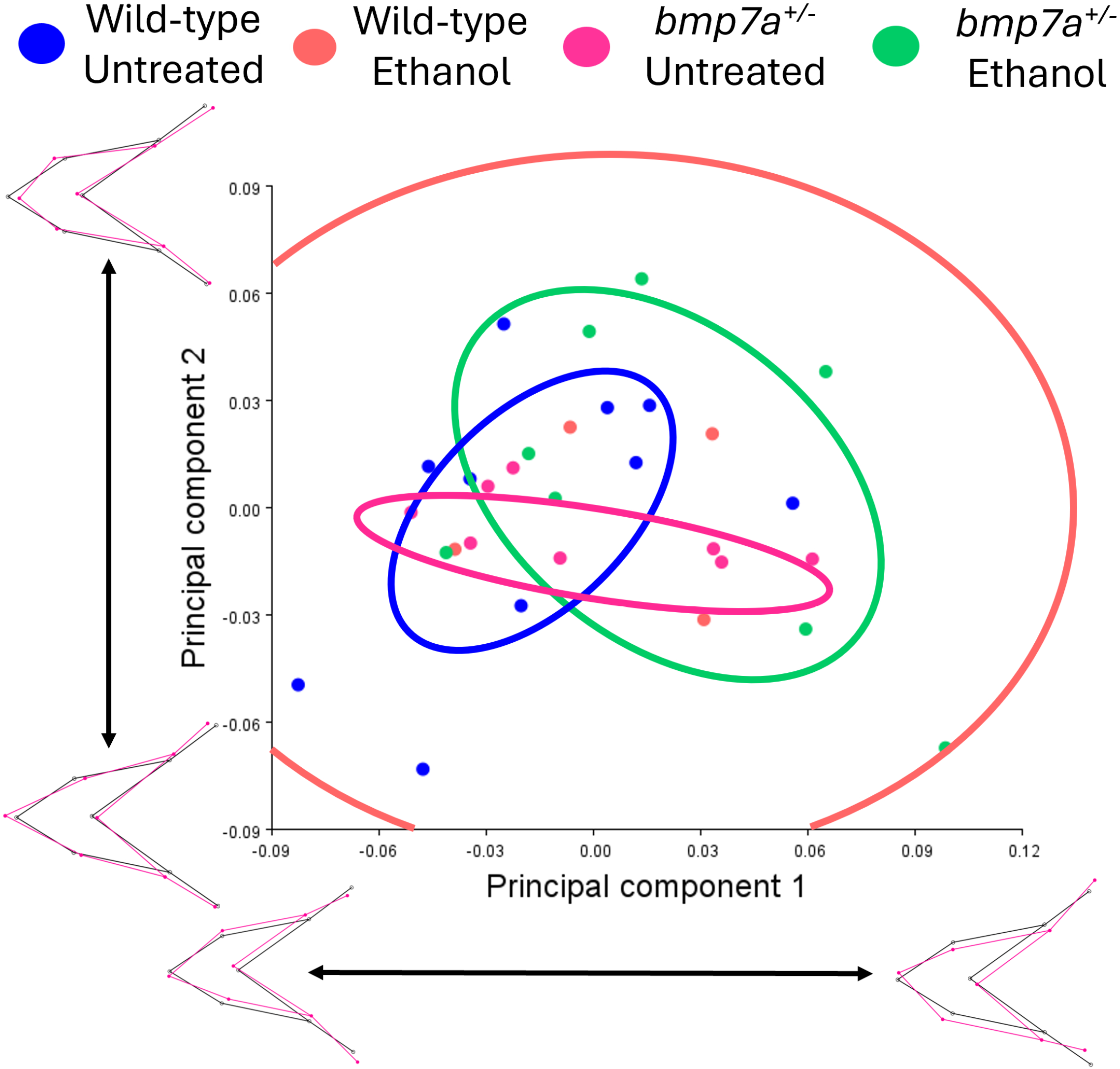
Principal Component Analysis plot of *bmp7a*^*+/-*^ and wild-type siblings with or without ethanol treatment, showing mean variation in facial shape. Genotypes are color-coded: blue = untreated wild-type larvae (n = 6), peach = ethanol-treated wild-type larvae (n = 4), magenta = untreated *bmp7a*^*+*/-^ larvae (n = 8), green = ethanol-treated *bmp7a*^*+*/-^ larvae (n = 8). Solid circles represent 95% confidence ellipses for the means. Wireframe graphs represent variation described by each PC axis, with black representing no variation and magenta representing variation relative to the black wireframe.

**Figure 3.**
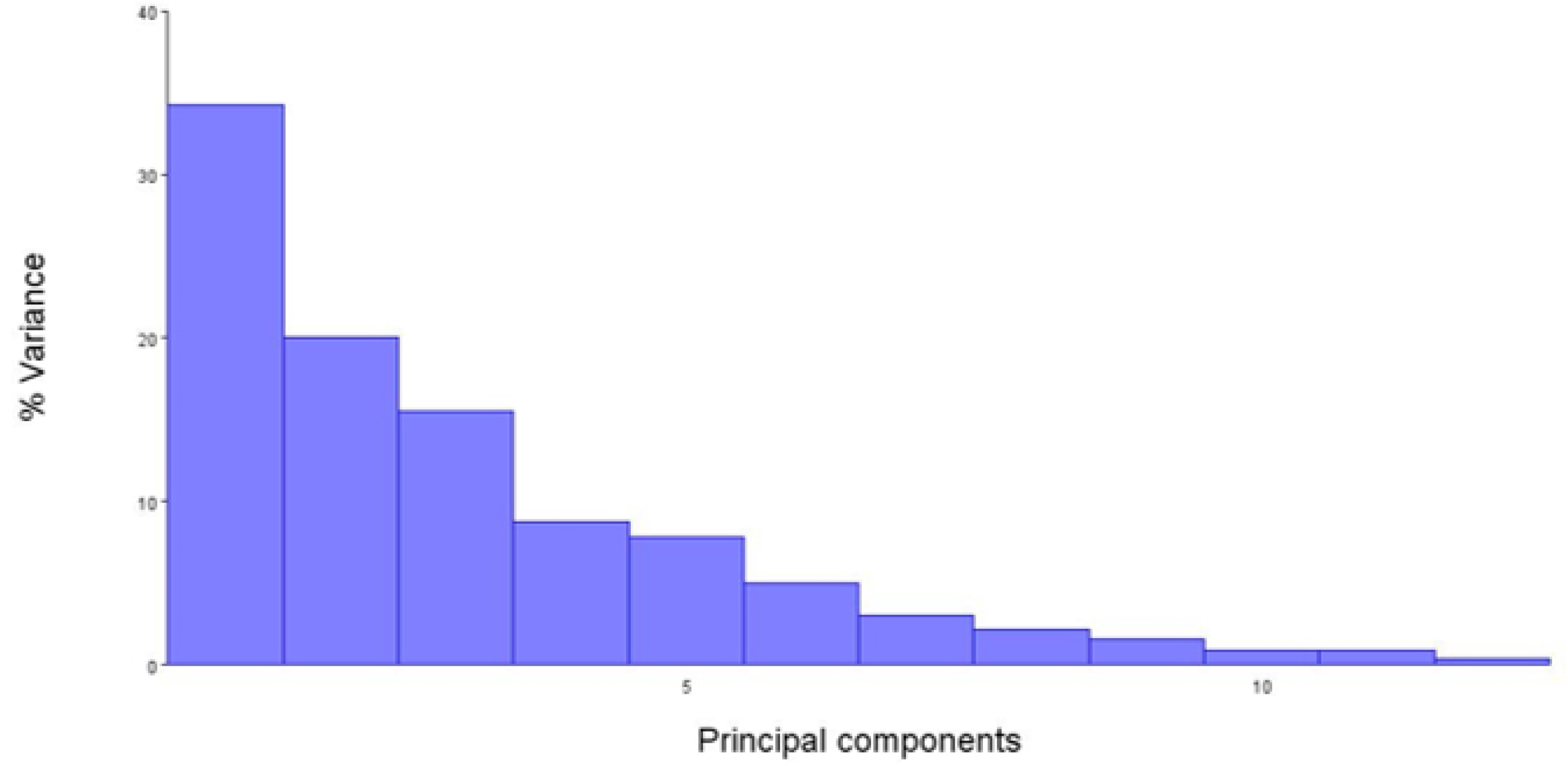
Scree plot showing the percentage of variance explained by each principal component in the PCA analysis in Figure 2. Over 50% of the variation in facial shape is defined by PC1 and PC2, with PC1 representing over 30% of the variation.

With solid ellipses represent the 95% confidence of the mean, the PCA plot shows overlapping means among the genotype and treatment groups, with no distinct clustering of the groups (Fig. 2). However, some of the means do spread along different PC axes, with both untreated and ethanol-treated *bmp7a* heterozygous larvae having greater spread along PC1, while untreated wild-type larvae spread more along PC2 (Fig. 2). The Ethanol-treated wild-type larvae exhibit a large confidence ellipse of the mean due to the low “n” of that group (Fig. 2). To test if this subtle variation observed in the PCA plot was significant, we performed a Procrustes ANOVA to assess variation in viscerocranial size and shape. No significant differences were detected in viscerocranial size (F = 1.47, p = 0.2445) or shape (F = 0.95, p = 0.5609; Pillai’s trace = 1.09, p = 0.7411), indicating that while varied, the overall size or geometric configuration was not significantly different in the dataset. This is consistent with the overlap observed in the means of each group and the lack of distinct clustering. However, the spread along different PC axes suggests that there might be some differences between the genotype and treatment groups that PCA and Procrustes ANOVA do not pick up.

A limitation of the Procrustes ANOVA in MorphoJ is that it does not analyze differences in specific facial shape changes, nor does it allow for pairwise comparisons of these differences between groups. To identify and analyze these differences, we used Canonical Variate Analysis (CVA). As described for PCA, CVA was conducted to visualize the distribution of individuals based on a specific shape variation, represented by a specific Canonical Variate ( CV1, CV2, etc.) (Fig. 4, solid ellipses represent the 95% confidence of the mean). The CVA plot shows that the means for untreated wild-type and untreated and ethanol-treated *bmp7a* heterozygous larvae do not overlap with each other (Fig. 4). Again, ethanol-treated wild-type exhibit a large confidence ellipse of the mean that overlaps all other groups due to the low “n” (Fig. 4). This approach generated raw CV scores, which we exported into PAST and performed a multivariate analysis of variance (MANOVA). The MANOVA revealed a significant overall effect of genotype and treatment (Table 1, Wilks’ λ = 0.2407, F = 5.176, p = 3.67e-05). Post hoc pairwise comparisons indicated a significant difference between untreated *bmp7a* heterozygous and wild-type larvae (p = 0.0033), with smaller effects observed for untreated *bmp7a* heterozygous larvae vs. ethanol-treated *bmp7a* heterozygous larvae (p = 0.0141) and untreated wild-type and ethanol-treated *bmp7a* heterozygous larvae (p = 0.0105).

**Table 1.** MANOVA results for untreated and ethanol-treated wild-type and *bmp7a*^*+/-*^ larvae. Results show significant differences between untreated vs. ethanol-treated wild-type and *bmp7a*^*+/-*^ larvae (Wilks’ lambda: 0.2407; F = 5.176; p = 3.67e-05).

| Table 1 |  |  |  |  |
| --- | --- | --- | --- | --- |
|  | Wild-type<br>Untreated | Wild-type<br>Ethanol | *bmp7a*<sup>+/-</sup><br>Untreated | *bmp7a*<sup>+/-</sup><br>Ethanol |
| Wild-type<br>Untreated |  |  |  |  |
| Wild-type<br>Ethanol | 0.21181 |  |  |  |
| *bmp7a*<sup>+/-</sup><br>Untreated | 0.0033455 | 0.066435 |  |  |
| *bmp7a*<sup>+/-</sup><br>Ethanol | 0.010515 | 0.073213 | 0.014133 |  |

**Figure 4.**
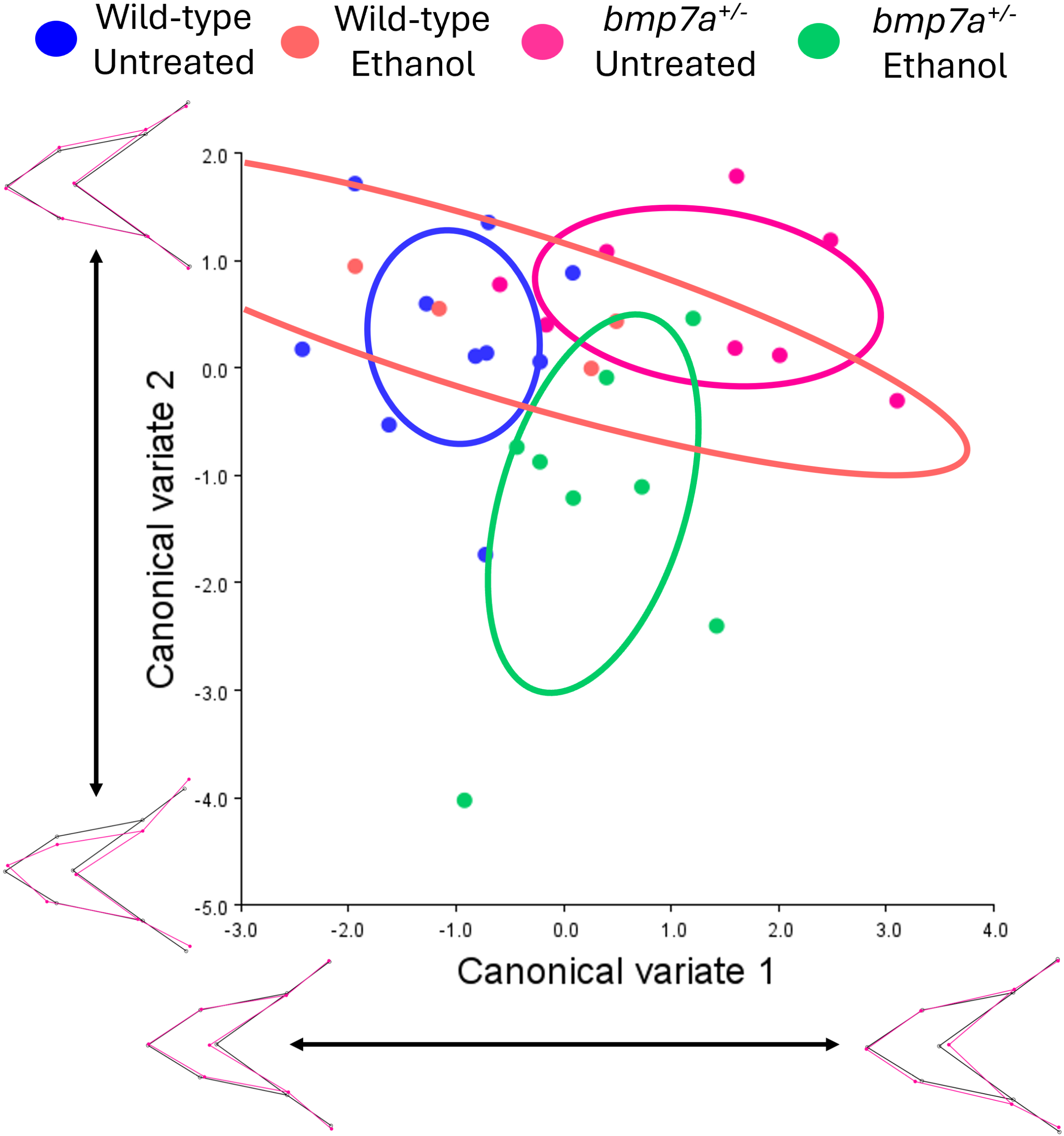
Canonical Variate Analysis plot of *bmp7a*^*+/-*^ and wild-type siblings with or without ethanol treatment, showing changes in specific facial shapes. Genotypes are color-coded: blue = untreated wild-type larvae (n = 6), peach = ethanol-treated wild-type larvae (n = 4), magenta = untreated *bmp7a*^*+*/-^ larvae (n = 8), green = ethanol-treated *bmp7a*^*+*/-^ larvae (n = 8). Solid circles represent 95% confidence ellipses for the means. Wireframe graphs represent variation described by each CV axis, with black representing no variation and magenta representing variation relative to the black wireframe.

## DISCUSSION

FASD is characterized by a wide range of developmental defects, including craniofacial defects such as jaw hypoplasia. Zebrafish are a strong model for FASD due to their genetic conservation with humans, high fecundity, translucent larvae, and external fertilization. Zebrafish have been used for decades to study both the formation of the craniofacial skeleton and the impact of ethanol on development [6, 12, 13, 15, 18, 27, 28]. More recent work has been exploring the impact of ethanol on facial development. Key to this work is the proper analysis of facial shape. In this paper, we have described a technique for analyzing facial shape holistically to determine significant morphological changes in jaw shape with FASD.

Quantifying facial shape is difficult for several reasons: linear measures of single facial structures are limited in quantifying overall morphological shape changes, impacts on overall growth confound measurable differences, and subtle differences that may not be noticeable to the eye. This is never clearer than in the work from Suttie et al. [29] that shows that classic linear measures of the cardinal facial features of FAS do not fully represent the broad impact of ethanol on facial shape. Using surface modeling and morphometric measures of shape, Suttie and colleagues show that ethanol induces a wide range of facial malformations beyond the cardinal measure of FAS, demonstrating the need to account for facial shape in totality in FAS. While PCA and additional analyses have been used more and more in recent FAS and FASD facial analysis in humans, these modalities have not been fully applied to animal models, wholesale [19]. Here, we show our approach using freeware morphometric and statistical software, TPSDigs2, MorphoJ, and PAST to generate PCA and CVA plots, and Procrustes ANOVA and MANOVA analyses examining facial shape in untreated and ethanol-treated zebrafish larvae.

We have previously shown that multiple ethanol-sensitive genetic loci contribute to a wide range of ethanol-induced malformations of the viscerocranium [6, 10, 18]. Our main focus of research as the ethanol-sensitive components of the Bone Morphogenetic Protein (Bmp signaling pathway. Bmp signaling is heavily implicated in craniofacial development, impacting multiple tissues in the developing head [30]. We show that mutations in *bmp2b, bmp4*, and *bmpr1bb* sensitize embryos to ethanol-induced facial dysmorphology by disrupting the formation of the pharyngeal endoderm, which leads to increased cell death in the cranial neural crest (which gives rise to the majority of the facial skeleton) [6, 10-12, 30, 31]. However, changes in embryos lacking one or both copies of *bmp7a* have not been directly measured, despite evidence supporting the role of *bmp7a* in facial development [32]. Embryos homozygous for mutation in *bmp7a* die during the gastrulation stage, so we assessed *bmp7a* heterozygous larvae and their wild-type siblings to quantify ethanol-induced facial shape changes [32]. Larvae were either treated with ethanol 6-24 hpf, which covers the time window of ethanol sensitivity of *bmp2b, bmp4*, and *bmpr1bb* [10].

To examine ethanol-induced changes to facial shape in wild-type and *bmp7a* larvae, we used software that is publicly available. TpsDig2 software allows for the preparation of the images to be analyzed by adding the landmarks to images that are needed for the morphometric analysis. TpsDig2 can also generate linear measures of structures in your images. MorphoJ creates a wireframe using landmarks assigned by TpsDig2 and visualizes shape change data for each sample compared to the average of all images analyzed. It does this by extracting shape data via Procrustes superimposition, which controls for size changes between images, maintaining all the necessary information for further analyses such as Procrustes ANOVA [20]. Though MorphoJ does not analyze differences in single facial shape changes or make pairwise comparisons between groups, it can generate CVs that can be exported to the statistical software package PAST [26]. PAST was originally designed for analyses in quantitative paleontology, but can be used with CV data from MorphoJ to generate pairwise comparisons of facial shape changes.

Here, we used this combination of software to show that ethanol does not impact the facial morphology of *bmp7a* heterozygous larvae compared to untreated and ethanol-treated wild-type siblings. The morphometric analyses through the Procrustes ANOVA indicated no differences in size (p = 0.2445) or shape (p = 0.5609), consistent with the PCA plots partially overlapping among groups due to lack of distinct clustering (Fig. 2). However, our sample size was small, with an “n” of 10 as the largest group. Our previous work used up to 58 larvae per group [10]. Thus, the small sample size may be decreasing our significance. Power analyses should be used to determine the appropriate “n” for each experiment. In contrast, a MANOVA revealed a significant overall effect of genotype and treatment on the combined dependent variables (Wilks’ λ = 0.2407, F = 5.176, p < 0.001). The most pronounced differences occurred between untreated *bmp7a* heterozygous and untreated wild-type larvae. However, CVA plots show the facial shape change driving these differences to be extremely small. While a larger “n” may increase the severity of shape differences, it is unlikely to be significant, as the results observed raise the question of the biological relevance of the statistical significance of the results. In other words, is the difference great enough to disrupt biological function?

While our approach is robust, there are a few key difficulties that can limit the utility of our approach. First, to use these software programs in tandem, the user requires a Windows-based computer system, as TpsDig2 and PAST software are not available to Mac- or Linux-based systems. Although this is an inconvenience, it can limit access due to the computer requirements of the programs. Second, while this methodology speeds up the assessment of facial shape changes, it can still be limited by the time required to mount and image the facial skeleton. In addition, improper positioning of the mounted larvae, leading to slightly tilted or uneven mounting, could account for disproportionate measurements, altering the dataset before morphometric analyses. This could be due to general developmental defects such as edema, which would impact the relative positioning of the facial skeleton. This inaccurate mounting, in combination with a low “n”, may be the reason for such a high variance result for untreated wild-type larvae. Third, our images are 2D, meaning we are missing a wealth of information from the 3^rd^ dimension. Although this is unlikely to cause issues with the TPSDigs2, MorphoJ, and PAST software, as they can handle and analyze 3D data, the lack of information 3^rd^ dimension could alter the interpretation of our results. With these points taken into consideration, our method successfully analyzes facial shape, resulting in a reliable quantification of facial morphology.

Zebrafish models of FASD have proven incredibly insightful in modeling ethanol-induced facial defects in humans [10, 33]. Through the morphometric approach described above, we have aligned facial shape changes observed in humans with those in zebrafish and identified genetic variants contributing to these facial defects [10]. This approach is incredibly versatile and can be applied to structures beyond the face, leading to new insights into form and function during development. Overall, this method will improve research in the structural impacts observed in FASD, and through the analysis of shape changes, can be used to quantify any number of structural malformations in multiple model systems.

## ACKNOWLEDGEMENTS

The research presented in this article was supported by a grant from the National Institutes of Health/National Institute on Alcohol Abuse (NIH/NIAAA) R01AA031043 to C.B.L.

## DISCLOSURES

The authors have nothing to disclose.

## Notes

### Competing Interest Statement

The authors have declared no competing interest.

